# Conformational locking of Ufm1 upon binding to the Ufm1-interacting sequence of Uba5

**DOI:** 10.1101/161802

**Authors:** Ryan T. Kelly, Walaa Oweis, Reuven Wiener, Christopher E. Berndsen

**Author notes:** To whom correspondence should be addressed: 901 Carrier Dr, MSC 4501, Harrisonburg, VA 22807.

## Abstract

Ubiquitin fold modifier 1 (Ufm1) is a ubiquitin-like protein (UBL) found in eukaryotic organisms which plays a crucial role in ER stress management and signal transduction. The crystal structure of UFM1 and its E1 (Uba5) in complex shows that Ufm1 binds to the adenylation domain of UBA5 and interacts with a separate Ufm1-interacting sequence (UIS) in the C-terminus of UBA5. The UIS interacts with Ufm1 on the opposite side of Ufm1 protein from the adenylation domain of Uba5 and the reason for this second interaction site is unclear. We analyzed Ufm1 bound to the UIS sequence through molecular dynamics simulations in order to identify additional functions for this interaction. We found that the residues in the adenylation interaction site of Ufm1 have less movement when the UIS peptide was bound to Ufm1 and formed a structure that aligns well with Ufm1 bound to the Uba5 adenylation domain. We further identified an amino acid that connects the UIS to the adenylation domain interacting site. Mutation of this amino acid decreases charging activity and shifts the Ufm1 conformation population toward the unlocked configuration even in the presence of the UIS peptide. These data suggest a role for the Uba5 UIS in stimulating activation of Ufm1.

## Introduction

Protein modification by ubiquitin or ubiquitin-like (Ubl) proteins regulate most cellular pathways ^1^. Ubiquitin and Ubl proteins share a common structure, the beta-grasp fold, despite little to no sequence similarity among the members of this family^1^. In addition to the differences in sequence, ubiquitin and each Ubl protein appear to have a distinct set of enzymes that transfer the Ubl protein to target proteins leading to distinct cellular functions^2–4^. While much is known about the structure of Ub, Ubls, and the conjugating enzymes, less is known about the chemical and regulatory mechanisms of the enzymes.

Ubiquitin or Ubl proteins are typically attached to the side chain of lysine, a process sometimes referred to as conjugation, using three distinct enzyme families^1^. The first enzyme, called the activating enzyme or E1 enzyme, adenylates the C-terminus cleaving ATP in the process. The Ub/Ubl-adenylate is then cleaved by a cysteine within the enzyme to form a thioester-linked Ub/Ubl∼E1 enzyme intermediate^2^. The second enzyme, called the conjugating enzyme or E2 enzyme, then binds to E1 and a transthioesterification reaction occurs transferring the Ub/Ubl protein to a cysteine within the E2^3^. Finally, the ligating enzyme or E3 enzyme, stimulates transfer of the Ub/Ubl protein to the substrate or a second transthioesterification reaction occurs to attach the Ub/Ubl protein to the E3 enzyme, which then modifies the protein substrate^4^. While there are several structures of enzymes in all three classes, the biochemical mechanisms of enzymes involved in Ub/Ubl transfer are still under investigation.

Ufm1 (Ubiquitin-fold modifier 1) is a Ubl protein found in metazoans linked to erythropoiesis and the ER stress response^5–7^. Recent structural and biochemical work on the activating enzyme for Ufm1, Uba5, indicated that this enzyme uses a unique Ubl protein binding mechanism^8–10^. In the “trans-binding” mechanism, Ufm1 interacts with both subunits of the Uba5 dimer: one interaction with the adenylation domain of one Uba5 subunit and one with the Ufm1-interacting sequence (UIS) within the C-terminus of the other Uba5 subunit^8^. Despite the similarity in activating enzyme structure and ubiquitin/Ubl protein structure, ubiquitin and most other Ubls do not share this binding mechanism^2,11^. Ubiquitin, NEDD8, and SUMO, bind directly to the adenylation domain and form extensive contacts with their respective enzymes^12–15^. In contrast, ATG7, the activating enzyme for the Ubl protein ATG8, uses a C-terminal extension to recruit ATG8 to the adenylation domain, a mechanism that is analogous to that of Uba5 and Ufm1, however ATG8 appears to bind *in cis*^11,16^. The reason for bipartite mechanism of Ubl binding is not clear for either system.

The interaction between Uba5 and Ufm1 is well described structurally however the necessity for the interaction with the UIS, beyond additional affinity, for the enzyme is unclear. Uba5 lacking the UIS cannot be activated, even at high concentrations of Ufm1, despite the larger interaction with the adenylation domain suggesting an additional role or roles. We explored the effect of the UIS interaction on the structure of Ufm1 in order to identify additional role(s) for the UIS interaction in the activation of Ufm1. We found that a peptide of the UIS can stimulate Ufm1 activation by truncated, inactive Uba5 suggesting UIS binding may induce allosteric changes in Ufm1. Molecular dynamics simulations show that the interaction with the UIS induces conformational locking of a loop that binds to the adenylation domain of Uba5 and enhances the correlative motions of Ufm1. Biochemical analysis of Ufm1 mutants identified a site within the core of Ufm1 that connects the UIS binding site to the adenylation domain binding site within Ufm1. Ultimately, we conclude that the binding of the Uba5 UIS to Ufm1 stabilizes an adenylation domain-binding competent version of Ufm1. The increases in protein rigidity enhance affinity of Ufm1 for Uba5 leading to increased activation of the protein.

## Methods

### Molecular Dynamics simulations

The human Ufm1 used in simulations was taken from Molecule C of PDB entry 5IAA and removing all other molecules and solvent^8^. For the bound simulation model, all protein and non-protein atoms from structure 5IAA were removed except for Molecule C and residues 333-346 of Uba5. Each simulation was equilibrated in YASARA in explicit solvent at 310 K, 0.9% NaCl, and pH 7.4, using the AMBER^14^ force field with an electrostatic cutoff of 8 Å, in a simulation cell with periodic boundaries extending 15 Å around the model. Molecular dynamics simulations were performed in YASARA using an AMBER14 force field for 0.5 to 1.2 microseconds. The apo and bound Ufm1 simulations were repeated 3 or 4 times to ensure reproducibility of the observations. Simulations were run with a timestep of 2.50 fs with the temperature adjusted using a Berendsen thermostat as described by Krieger et al ^17^. Simulation trajectories were analyzed to produce data on the RMSD, RMSF, DCCM, and structures using the pre-packaged macros in YASARA. Cavity volumes were determined by counting the number of water molecules within 4 Å of Leu33 and calculating the volume of those water molecules.

### Protein expression and purification

Uba5^57-363^ was expressed in BL21(DE3)pLysS cells in 2xYT overnight at 16 °C. Ufm1 and Ufm1 mutants were expressed in BL21(DE3)pLysS cells in 2xYT expressed for 4 hours at 37 °C. Cells containing Uba5^57-363^ were lysed by sonication in 20 mM Tris, pH 7.5, 500 mM NaCl, 2 mM imidazole. Soluble lysate was separated from insoluble material by centrifugation at 17500 × g for 20 minutes and 4 °C. Protein was purified by high density cobalt resin (Gold Bio). Protein was washed in 20 mM Tris, pH 7.5, 500 mM NaCl, 10 mM imidazole buffer and eluted with 20 mM Tris, pH 7.5, 50 mM NaCl, 1 mM Tris(2-carboxyethyl) phosphine, 5 mM MgCl2, 200 mM imidazole buffer. Protein was then dialyzed (SnakeSkin 7,000 MWCO) into 20 mM Tris, pH 7.5, 40 mM NaCl 0.1 mM Tris(2-carboxyethyl) phosphine. Ufm1 and Ufm1 mutants were purified following the protocol of Oweis and coworkers^8^.

### Pulldown assays

Pulldown assays between Ufm1 and Uba5 or mutants of Uba5 were performed largely as described previously in Oweis, et al., 2016 except 100 µg of GST-Uba5^314-404^ was used in place of full length Uba5. All other steps were performed as described^8^.

### Ufm1 Charging Assays

Charging assays were performed using Uba5^57-363^ as described previously by Oweis, et al., 2016^8^. Briefly, reactions contained 20 µM Uba5, 100 µM Ufm1 or Ufm1 mutants, 500 µM ATP in 50 mM MES, pH 6.5, 100 mM NaCl, 10 mM MgCl_2_. V66A reactions were performed with 50 µM Ufm1. Reactions were incubated at 30 °C for the times incubated before being quenched in non-reducing SDS-PAGE loading dye. Samples were separated on MOPS 4-20% gradient gel and stained with Coomassie Brilliant Blue.

For charging assays of Uba5^57-329^ with the UIS peptide, Uba5 (57-329) at 20 μM concentration was incubated with different concentrations of the UIS peptide (IIHEDNEWGIELV), or 1mM of the scrambled peptide (ILDIWHEIGNEEV) at 30°C for 90 min in 50 mM Tris–HCl pH 7.5, 150 mM NaCl, 5 mM MgCl_2_, 5 mM ATP.

### Circular Dichroism

Data on Ufm1 wild-type or mutants were collected in 20 mM phosphate (pH 7.5) in a 2 mm quartz cuvette. Spectra were recorded using a JASCO J-810 spectropolarimeter from 190 nm to 300 nm with a 0.2 nm step size at a rate of 100 nm/min. Each spectrum is an average of three accumulations from 190 to 290 nm.

## Results

### Activation of Uba5^57-329^ by UIS peptide

Previously, we and others characterized the trans-binding mechanism of Uba5 for interacting with Ufm1 and described how a C-terminal sequence is required for high affinity binding and activation of Ufm1^8–10^. While it is clear that this region is important for binding to Ufm1 and luring it to the adenylation domain, we speculated additional functions. Therefore, we tested whether a peptide of the UIS (Uba5 amino acids 334 to 346) could be added *in trans* and stimulate activation of an inactive truncation of the E1, Uba5^57-329^. Addition of the peptide increased the amount of thioesterified Ufm1 bound to Uba5 (Figure 1). Use of a peptide with the same amino acid content, but with a distinct order from the UIS peptide or prevention of thioester formation by using a C250A mutation within Uba5, decreased activity. These data indicate that the increase in Ufm1∼Uba5 shown in the gel in the presence of the UIS peptide is specific and is the formation of a thioester and not tight, non-covalent binding of Ufm1 to Uba5. The results of this experiment show that the UIS peptide influences Uba5 activity in an additional way to serving as a binding domain within Uba5.

**Figure 1.**
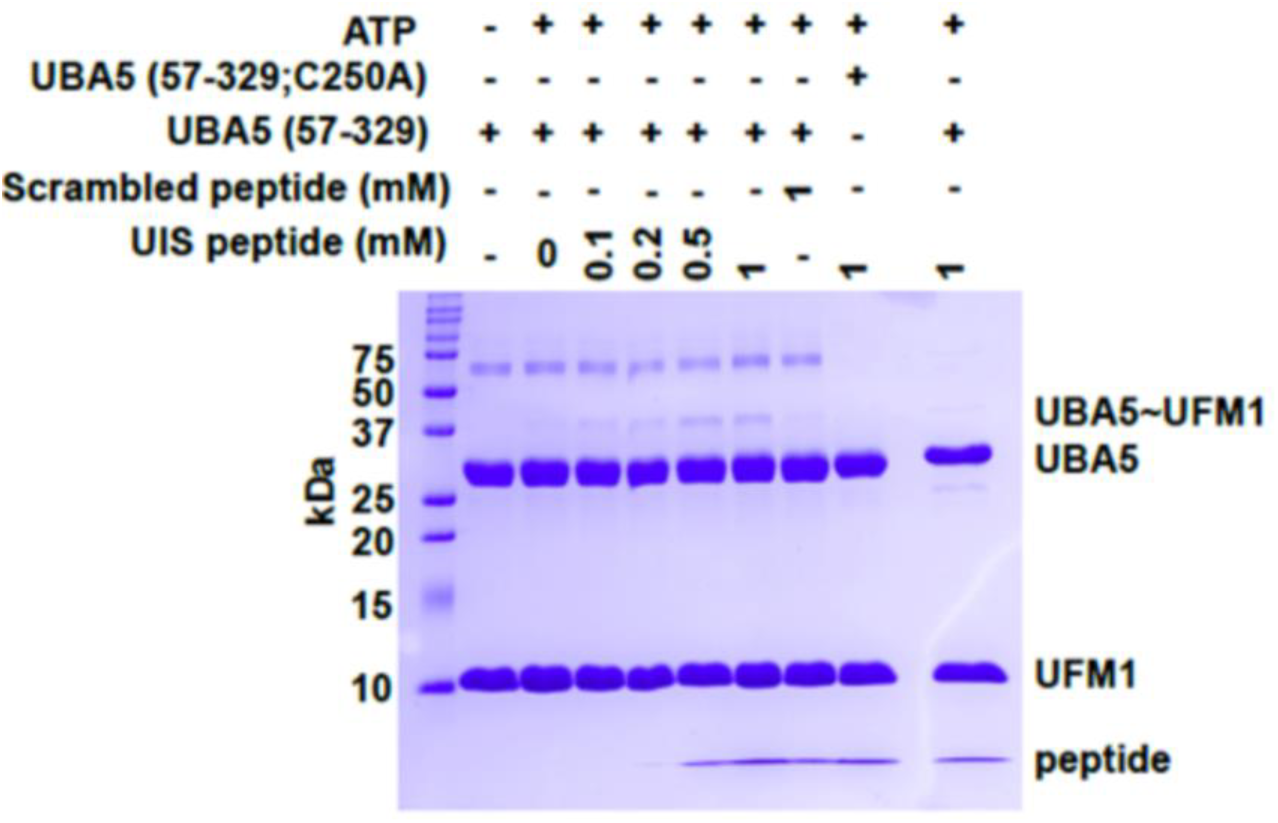
Charging of Uba5 is stimulated by a peptide of the UIS. Charging of Uba5^57-329^ in the the absence or presence of increasing concentrations of a UIS peptide (Uba5 residues 334 to 346) or a peptide with the scrambled sequence of the UIS. Catalytically inactive UBA5 C250A was used as a negative control.

### UIS interaction with Ufm1 restricts the dynamics of Ufm1

The finding that the UIS stimulates Uba5 activity when added *in trans* suggested structural changes in Ufm1 may be occurring, despite crystallographic and NMR studies of Ufm1 bound to Uba5 or the UIS not showing significant changes in structure^8–10^. However, NMR solution structures of human or mouse Ufm1 alone suggest several dynamic regions indicating that the UIS may alter the motions of Ufm1 rather than inducing large scale structural changes^18,19^. To observe possible changes in dynamics, we performed molecular dynamic simulations of Ufm1 alone and of Ufm1 in the presence of amino acids 333-346 of the UIS. The duration of each simulation was at least 1 µs and performed twice for each condition. General conclusions are drawn from cumulative observations of the datasets. Initially, we compared the root mean square deviation (RMSD) for structures from every 10 ns to the energy minimized structures from both simulations. While the apo and bound conditions showed similar changes in RMSD value over time, the distribution of the data between the two simulations was distinct (Figure 2A). Comparison of the RMSD of simulation snapshots to the crystal structures of the bound and unbound structures of Ufm1 revealed that Ufm1 bound to the UIS clustered in a smaller range of RMSD values compared to the apo simulation data, suggesting a decrease in the dynamics of Ufm1 upon binding to the UIS (Figure 2B).

**Figure 2.**
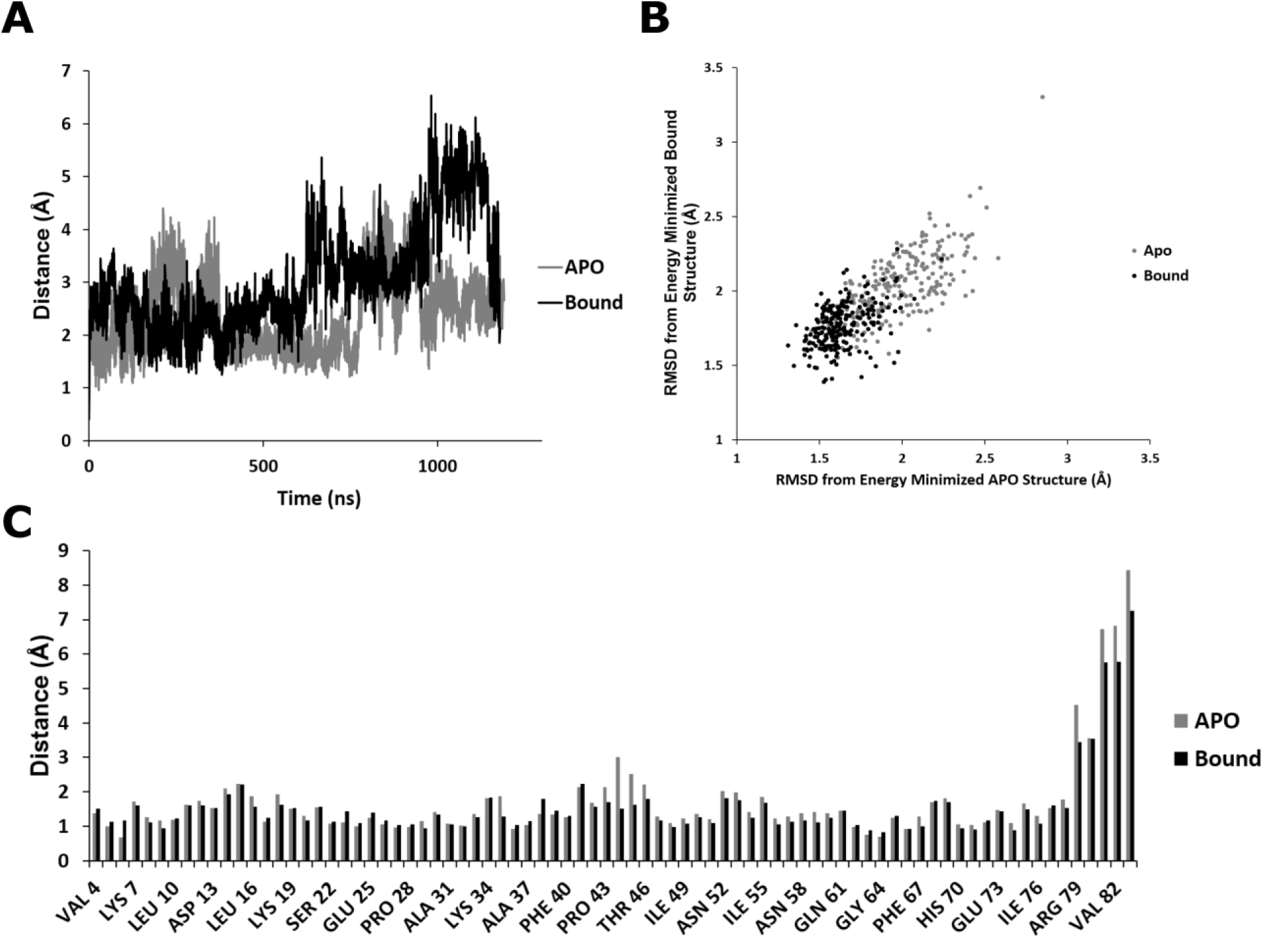
Simulations of UFM1 APO and bound to UIS. **(A)** RMSD of the APO (grey) and bound (black) simulation at 310K. **(B)** The RMSD of structures every 10ns in the APO and bound simulations compared against the energy minimized APO and bound structures. **(C)** The RMSF of each residue throughout both simulations.

In order to locate the specific amino acids which were affected, we measured the root mean square fluctuation (RMSF) value for the alpha carbons of each amino acid, which shows how mobile each alpha carbon is during the simulation. We did not include the first 100 ns of the simulation in our analysis to remove effects of the protein adapting to the forcefield. Comparison of the per residue RMSF values, shows that overall the dynamics of Ufm1 are comparable except for the C-terminus, which is known to be flexible, and amino acids 43 to 46, which showed higher RMSF values in the apo simulation (Figure 2C). This region of the protein is on the opposite face of the Ufm1 from the UIS interaction site. Alignment of snapshots taken every 25 ns from the apo simulation and bound simulations show that in the apo simulation the loop containing these amino acids appears to adopt two distinct positions, while the loop position is largely static in the bound simulations (Figure 3A). The mobile loop is part of a larger structure that is known to interact with Leu233 within the adenylation domain of Uba5 (Figure 3B)^8^. We next measured the distance between the alpha carbon of Ala45 and the alpha carbon of Pro59 every 0.25 ns during the simulation. These amino acids are the tips of the region which binds to Leu233 in the crystal structure (Figure 3A). Plotting these measurements in 0.2 Å bins as a histogram shows that in the bound simulations the loop adopts a structure such that the distance is near the value observed in the crystal structure 5IAA (Figure 3C)^8^. In the apo simulations, there are two distinct populations, neither of which corresponds exactly to the structure observed in the bound simulations or the crystal structures (Figure 3C). We further measured the volume of the cavity adjacent to this loop finding that the distribution of volumes was smaller in the bound simulations than those values measured from the apo simulations (Figure 3D). Alignment of Ufm1 highlights the adenylation domain interacting site (referred to as the AIS for the rest of the manuscript) in the open and closed conformations, and shows that the UIS bound structure matches the crystal structure in this loop region, while the apo structure does not (Figure 3B)^8^. Cumulatively, these data suggest that the bound simulation maintains the Ufm1 structure such that it is in the conformation that binds the adenylation domain more favorably.

**Figure 3.**
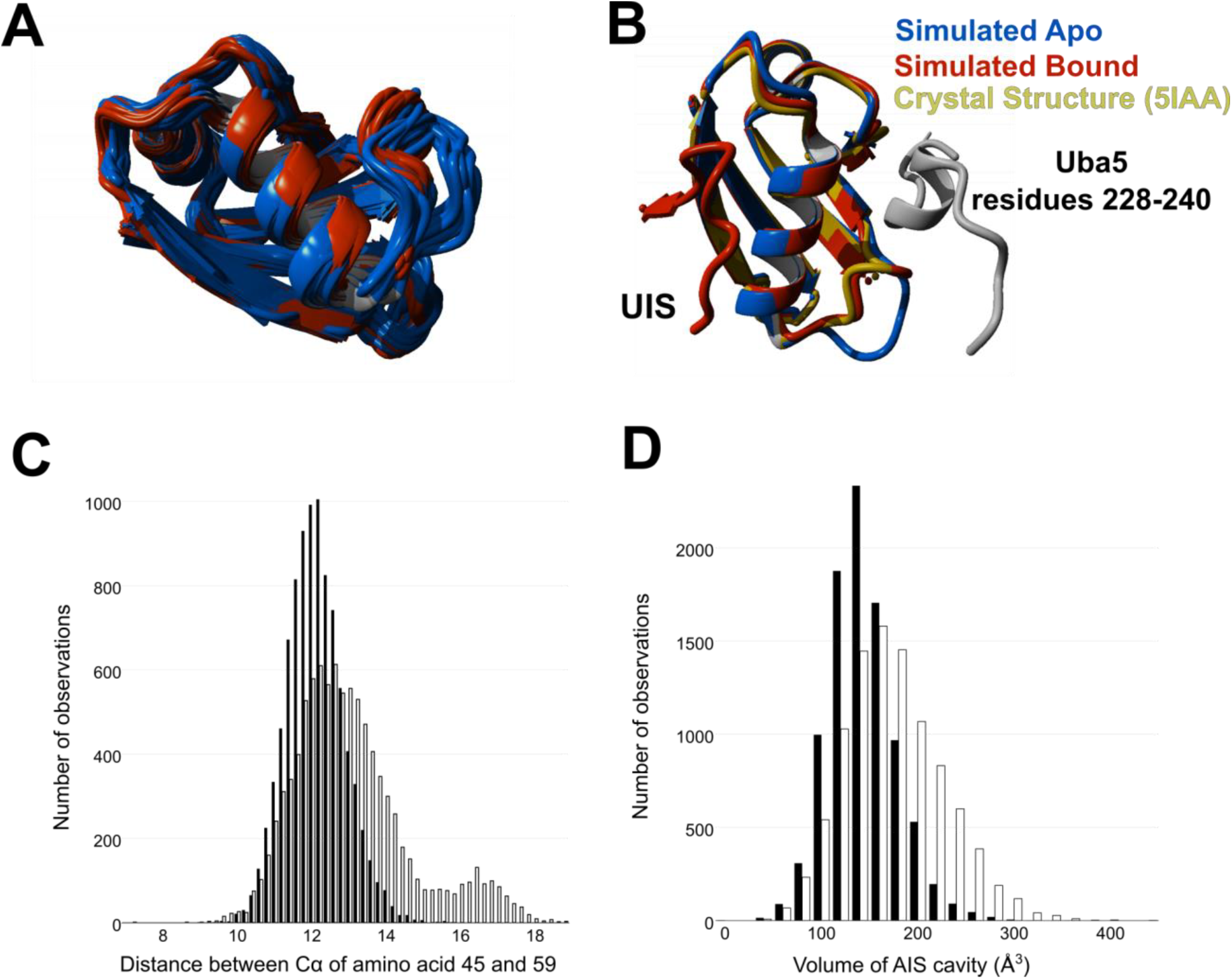
Structural Deviations Between Apo and Bound Simulations. **(A)** Aligned simulation snapshots taken every 25 ns from the APO simulation (blue) and bound simulation (red). **(B)** An APO simulation snapshot with an open AIS conformation (blue) and a bound simulation snapshot in the locked AIS conformation (red) compared to the UFM1 crystal structure (5IAA, Mol C, yellow) and an interacting helix from the E1 (5IAA, Mol A, grey), however was not included in the simulation. **(C)** Distance between Ala 45 and Pro 59 every 25ps over the course of both 310K simulation for the APO (white) and bound (black) simulations. **(D)** The volume of the AIS cavity in the APO (white) and bound (black) simulations.

### UIS increases correlative motions within Ufm1

The changes in the structure of the mobile loop within the adenylation domain interacting site upon binding the UIS led us to try to identify the structural features of Ufm1 that produce the overall increase in stability. To identify the network of amino acids that controlled the structural changes, we compared the dynamic cross-correlation matrices for each simulation. In the bound simulations, we observe higher correlations between amino acids, suggesting tighter packing of the amino acids (Figure 4A). In contrast, the apo simulation shows higher levels of anti-correlation between amino acids (Figure 4B). These data match our RMSF data showing less motion in the bound simulations and indicating that the UIS induces Ufm1 to form a more rigid structure.

**Figure 4.**
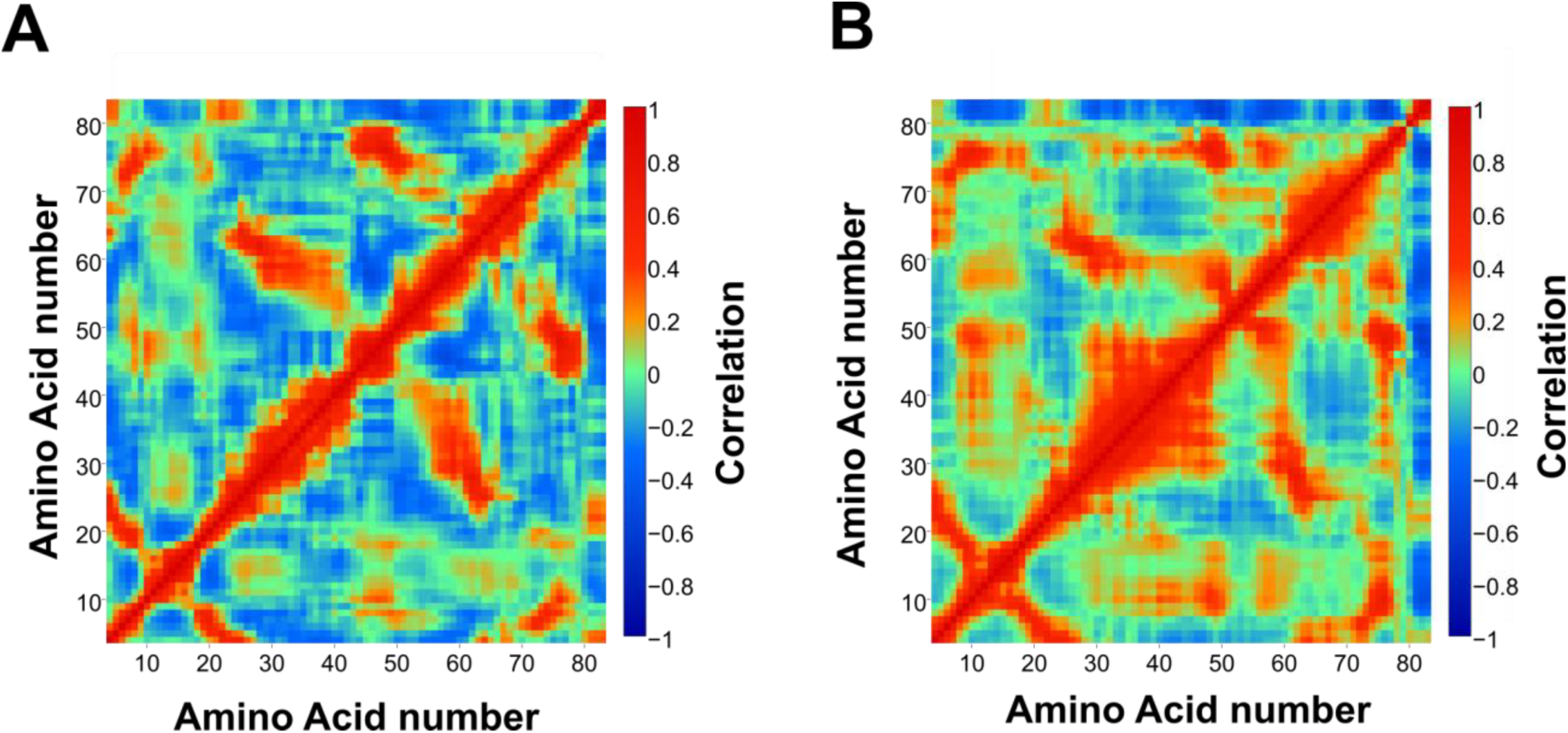
Correlation Between Residues in Both Simulations. **(A)** Dynamic cross-correlation matrix (DCCM) for the APO simulation shows the correlation in fluctuations for each residue as compared to every other residue. **(B)** DCCM for the bound simulation excluding correlations comparing residues in the UIS. Red coloring indicates higher correlation of motion, while blue coloring indicates anti-correlative motions.

### Mutations in AIS decrease rate of Ufm1 charging by Uba5

To further explore this functions of the AIS, we made four mutations in the AIS and determined the effects of these amino acid substitutions on UIS binding and the charging of Uba5 with Ufm1. Mutation sites were chosen based on sequence conservation and proximity to the AIS and UIS interacting site. Mutations did not affect the secondary structure of Ufm1 proteins in circular dichroism assays (Supplemental Data figure 2A), however the mutations at V66A and L33A did cause increased precipitation of these proteins over time suggesting a decrease in stability (data not shown). None of the mutants had an effect on binding to the UIS in pulldown assays suggesting that the UIS and AIS sites are structurally independent (Supplemental Data Figure 2B). In charging assays, all mutant proteins affected charging relative to the wild-type reactions, however to different extents (Figure 5B). Leucine 33, which makes direct contact with Uba5, reduced activity to a low level, when mutated to alanine. Substitution of alanine for Threonine 46 or Serine 47, which are in the mobile loop and do not make direct contact with Uba5, reduce activity to roughly two-third to one half of the activity seen with the wild-type protein. Substitution of alanine for Valine 66, which contacts amino acids in the both the UIS and the AIS, reduced activity to a level comparable to that of Ufm1 containing the Leu33 mutation. Substitutions do not affect the stability of the Ufm1∼Uba5 thioester, as at longer time points, all mutations were charged to approximately the same degree (Supplemental Data Figure 2C).

**Figure 5.**
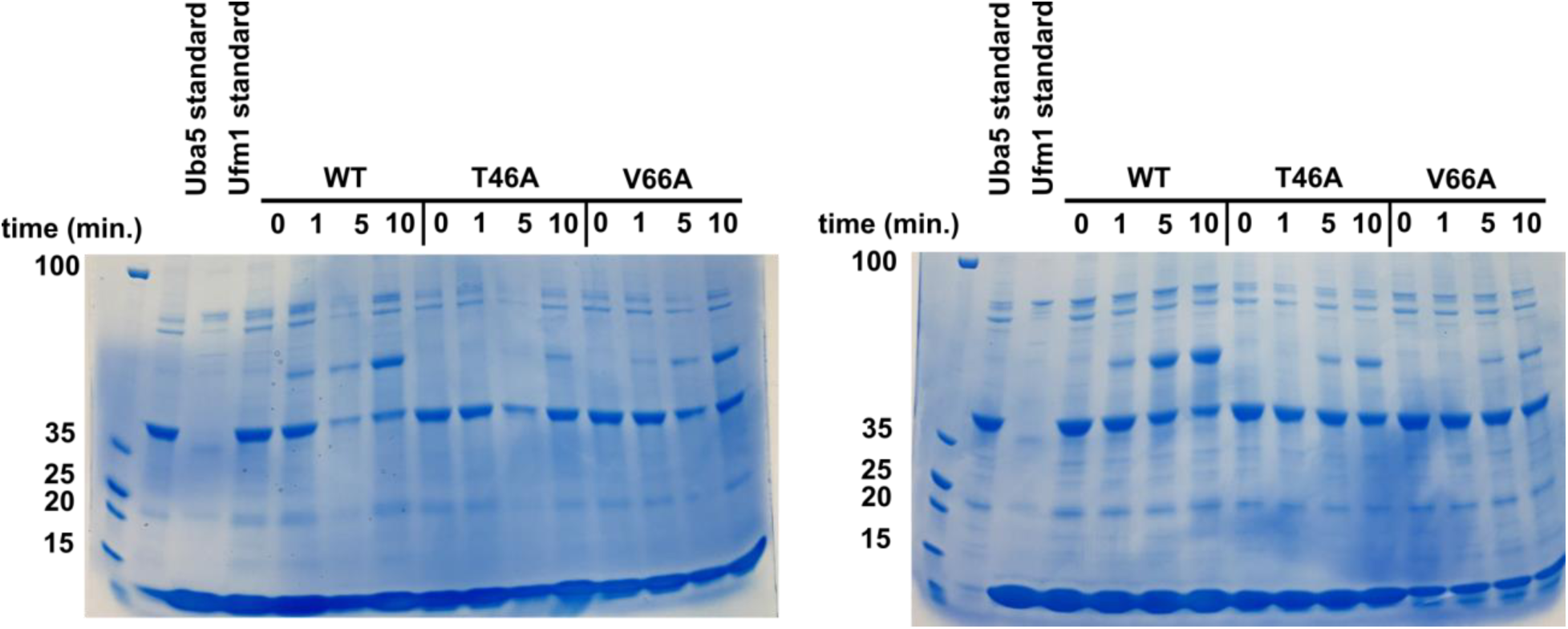
Charging of Mutants in the AIS. **(A)** Charging of UFM1 mutants L33A and S47A compared to wildtype. **(B)** Charging of UFM1 mutants T46A and V66A compared to wildtype. V66A is at half concentration compared to other UFM1 proteins.

We next did simulations of Ufm1 containing a S47A or V66A bound to the UIS peptide to illustrate the molecular basis for the activity defects. For both mutants, Ufm1 retained the overall structure during the simulations with RMSD values of 1.2 and 1.1 between the Ufm1 wild type in complex and the S47A and V66A Ufm1 proteins in complex, respectively (Figure XX). Comparison of the dynamic cross correlation matrices from the mutant simulations to the simulations of the UIS bound and unbound wild-type shows that the mutant matrices resemble the matrix for the bound WT simulation (Figures 6A and 6B). There are differences in anti-correlated motion the first 50 amino acids and amino acids 50 to 75 which indicates a loss of connection between amino acids in this region. Histograms of the distance between Ala45 and Pro59 for the S47A and V66A Ufm1 mutations show a distribution of distances that is intermediate between the wild-type bound and apo simulations (Figure 5C). The S47A mutation shifts the major peak center to longer distances by 0.2 Å but overall the distances fall largely within that of the WT bound simulation. The sidechain of Ser47 makes contacts with the Ufm1 backbone which may alter the AIS loop structure. The V66A mutation shows an increased number of structures with distances >16 Å which is in the region of the lesser peak of the apo Ufm1 simulation. These data suggest that the V66A mutation uncouples the UIS interacting site and the AIS within Ufm1 leading to defects in binding of Ufm1 to the adenylation domain of Uba5 and the activity defects observed in our assays.

**Figure 6.**
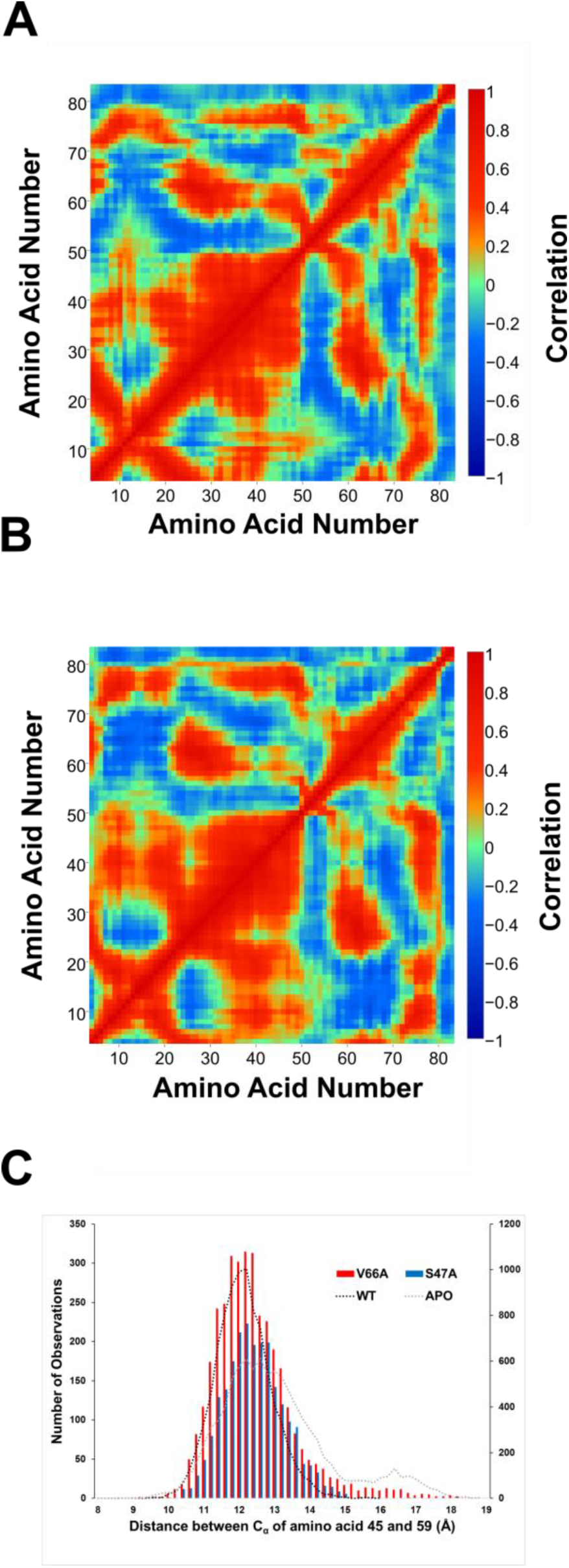
Correlation of Residues in Mutant UFM1 Bound Simulations. **(A)** DCCM for the mutant UFM1 S47A bound to the UIS. **(B)** DCCM for the mutant UFM1 V66A bound to the UIS. **(C)** Distance between Ala 45 and Pro 59 every 25ps over the course of both 310K simulation for the S47A (blue) and V66A (red) simulations. The data for the Bound (black dashed line) and the Apo simulations (grey dashed line) are from Figure 3.

## Discussion

The recently identified trans-binding mechanism of Ufm1 binding by the E1 protein Uba5 and the finding that a peptide of the Ufm1-interacting sequence could stimulate the activity of an inactive truncation of Uba5 led us to investigate the basis for the stimulation. Microsecond simulations of Ufm1 and Ufm1 bound to the UIS peptide revealed changes in the dynamics of Ufm1 upon peptide binding. These changes were most prevalent in a section of Ufm1 shown to bind the adenylation domain of Uba5. Mutations in Ufm1 that directly or indirectly appear to function in the connection between the UIS and AIS, showed no change in UIS binding, but did affect charging of Uba5. Simulations of the Ufm1 containing one of two of the mutations bound to the UIS show increased anti-correlated motions, indicating the mutations break the connection between the two regions. The V66A mutation in particular appears to increase the apo-like conformation of Ufm1 based on comparison of the Ala45-Pro59 distance histograms (Figures 3C and 6C). These data suggest V66 is part of the network of amino acids that connect the UIS and AIS sites on Ufm1. This connection allows UIS binding to configure Ufm1 to bind the adenylation domain of Uba5 leading to activation of the protein. Binding to the UIS of Uba5 restricts the dynamics of the loop so that the AIS is configured to effectively bind to the adenylation domain of Uba5.

Previous study of the interaction between Ufm1 and the Uba5 UIS by NMR did not identify allosteric or structural changes upon complex formation^10^. We believe this is because the structural changes are in fast exchange relative to the time scale of these previous NMR experiments^20^. Upon Ufm1 binding to the UIS, Habisov and coworkers did record chemical shift changes in some of the AIS residues including L33, A44, S47, and N58 however, with the exception of L33, these changes were one standard deviation above the average and therefore not considered significant by the authors^10^. In fast exchange for a two-state system, the chemical shifts will reflect the population weighted average of the states^20,21^. In our data, the AIS in the unbound simulations favor conformations near to or overlapping with the conformation of the bound ∼93% of the time based on the number of structures that have Ala45-Pro59 distances less than 16 Å (Figure 3C). In contrast, the V66A mutation is in the correct orientation ∼98% percent of the time (Figure 6C). Thus, binding the UIS shifts the population of Ufm1 conformations toward the complex state. However, given that the overall population spends part of the time in this conformation already, the chemical shifts are expected to be small in experiments that are not designed to measure dynamics and fast time-scale motions. Therefore, we believe our current findings provide further insight into the NMR experiments of Habisov and coworkers by showing distal changes in Ufm1 structure by binding to the Uba5 UIS.

Allostery and dynamics in ubiquitin and ubiquitin-like structure are a new area in the field. Recent picosecond to millisecond molecular dynamics simulations of ubiquitin revealed that ubiquitin by itself adopted a variety of conformations similar to that of ubiquitin bound to other partners^22^. Additionally, there were some correlation of motion across the protein. In this study, which sampled Ufm1 structure on the microsecond timescale, we too observed that Ufm1 visited conformations similar to Ufm1 bound to Uba5 (the only known ligand with a structure) and that there were correlated motions across the protein (Figures 3 and 4). We propose that this allostery facilitates Ufm1 binding to Uba5 leading to activation of the Ubl protein. Given the dynamics data on ubiquitin from Lindorff-Larsen and coworkers, it is possible similar mechanisms exist to enhance ubiquitin or Ubl proteins affinity for different proteins, however this remains to be fully demonstrated. Given the sequence differences between Ufm1, ubiquitin and other Ubl proteins, these motions may be quite dissimilar to those we propose for Ufm1.

## Acknowledgements

We thank Laurel Feichtel, Brett Siegel, and Sam Holmes for comments on the manuscript prior to submission. Dr. Linette Watkins for support and Dr. Isaiah Sumner for technical suggestions. The work was supported by a United States-Israel Binational Science Foundation Grant 2013261, a National Science Foundation Research Experience for Undergraduates Grant CHE-1461175, and the JMU Department of Chemistry and Biochemistry.

